# Cockroach bacteriocytes migrate into the ovaries for vertical transmission of the bacterial endosymbiont *Blattabacterium*

**DOI:** 10.1101/2025.06.22.660904

**Authors:** Tomohito Noda, Toshiyuki Harumoto, Tatsuya Katsuno, Minoru Moriyama, Takema Fukatsu

**Affiliations:** Department of Biological Sciences, Graduate School of Science, The University of Tokyo, Tokyo, Japan; Molecular Biosystems Research Institute, National Institute of Advanced Industrial Science and Technology (AIST), Tsukuba, Japan; Hakubi Center for Advanced Research, Kyoto University, Kyoto, Japan; Graduate School of Biostudies, Kyoto University, Kyoto, Japan; Life Science Center for Survival Dynamics, Tsukuba Advanced Research Alliance (TARA), University of Tsukuba, Tsukuba, Japan; Center for Anatomical Studies, Graduate School of Medicine, Kyoto University, Kyoto, Japan; Graduate School of Life and Environmental Sciences, University of Tsukuba, Tsukuba, Japan

**Keywords:** Cockroach, *Blattella germanica*, vertical transmission, bacteriocyte, *Blattabacterium*, symbiotic bacteria, insect ovary development

## Abstract

Diverse insect groups are obligatorily associated with and dependent on specific microorganisms as essential mutualistic partners that are usually maintained in specialized cells or organs, called bacteriocytes or symbiotic organs. Many organisms with symbiotic microorganisms have developed elaborate vertical transmission mechanisms, which are thought to be important for the evolution of intimate symbiotic relationships with microorganisms. One such case is the cockroach-*Blattabacterium* endosymbiosis, in which the symbiotic bacteria have been evolutionarily conserved and co-speciated with the host insects with stable vertical symbiont transmission via ovarial passage. While classical histological descriptions and recent electron microscopic observations have reported the vertical symbiont transmission processes in some cockroach-*Blattabacterium* associations, the full picture of the infection dynamics has not been fully understood. In this study, we conducted detailed histological and cytological observations of the localization of the bacteriocytes and the symbiotic bacteria during the postembryonic development of the German cockroach *Blattella germanica*. We found that the symbiont-filled bacteriocytes migrate into and associate with nymphal ovaries and are subsequently eliminated from adult ovaries, suggesting that symbiont infection to the ovaries may only occur during nymphal stages. We also found that the symbiotic bacteria are localized in the space between each oocyte and surrounding follicle cells, the symbiont-localized space is interconnected between neighboring oocytes, and therefore the symbiotic bacteria can move across oocytes within the same ovariole, suggesting the possibility that the symbiont-infected oocytes may serve as the source of symbiont supply to developing young oocytes upstream in the same ovariole. Based on these observations, we provide a hypothesis as to how the postembryonic developmental dynamics of the bacteriocytes are integrated into the vertical symbiont transmission and functioning in the cockroach-*Blattabacterium* endosymbiosis.

## Introduction

Diverse insect groups are intimately associated with microbial endosymbionts, many of which play significant biological roles for the host insects [1, 2]. These range from facultative symbionts that are not essential for survival and reproduction of the host insects, which may be either parasitic, almost neutral or conditionally beneficial as represented by *Wolbachia* [3, 4], to obligatory symbionts that are indispensable for survival and reproduction of the host insects such as *Buchnera*, the bacterial intracellular endosymbiont of aphids [5, 6]. Such obligatory microbial symbionts may supply essential nutrients [6, 7], assist food digestion [8, 9], defend against natural enemies [10, 11], etc., for the host insects. These obligatory microbial symbionts are of great interest not only in terms of the evolution of microbial symbiosis, but also from the perspective of insect diversity, as the biological functions provided by the symbionts have allowed the host insects to conquer novel ecological niches, thereby significantly contributing to the insect diversity we observe today [12].

In these symbiotic associations, the host insects have often evolved specialized cells, tissues, and organs for harboring the symbiotic bacteria. Some insects maintain their symbiotic bacteria extracellularly within the inner cavity of specialized structures associated with their alimentary tract called crypts or gastric caeca, as in stinkbugs and leaf beetles [13–15]. Other insects harbor their symbiotic bacteria intracellularly within the cytoplasm of specialized cells and organs, called bacteriocytes and bacteriomes, as in aphids and grain beetles [16–19]. The endosymbiotic bacteria of cockroaches, *Blattabacterium* spp., which are thought to play a role in the recycling of host’s nitrogenous waste products [20, 21], are harbored in the cytoplasm of specialized host cells called bacteriocytes [22, 23]. In most insects harboring obligatory endosymbiotic bacteria, the bacteriocytes tend to aggregate to form symbiotic organs called the bacteriomes [24–26]. In cockroaches, by contrast, the bacteriocytes do not form such bacteriomes. Instead, individual bacteriocytes are scattered within the abdominal fat bodies, exhibiting a unique bacteriocyte distribution among diverse insects [1, 23, 27]. *Blattabacterium* has been found in almost all known cockroach species, and its bacteriocyte distribution is highly conserved among them, suggesting that the cockroach-*Blattabacterium* endosymbiosis is of ancient origin and has been stably maintained over evolutionary time [20, 28, 29].

For ensuring stable maintenance of host-symbiont association and facilitating the evolution of mutualism, special mechanisms for vertical symbiont transmission through host generations have evolved in diverse insects, which are represented by smearing of symbiont-containing secretion onto the egg surface [30–32], deposition of symbiont-containing packets with eggs [33–35], direct symbiont transmission from bacteriocytes to early oocytes/embryos in the maternal body [16, 36], symbiont infection to oocytes via maternal germline cells [37, 38] and many others [1, 39]. As for the cockroach-*Blattabacterium* endosymbiosis, a number of classical microscopic descriptions and more recent cytochemical and electron microscopic observations of the symbiotic bacteria, the bacteriocytes and the vertical transmission processes have been reported [40–57]. Being one of the earliest described insect-bacterium endosymbiotic associations [40] along with the easy accessibility as a home-dwelling pest, the cockroach-*Blattabacterium* association represented one of the best-studied insect-microbe symbiotic systems in the 20th century. Although more recent molecular phylogenetic studies [58–64] and comparative symbiont genomics [20, 28, 65–71] have been compiled, many biological and functional aspects of the cockroach-*Blattabacterium* associations have been still elusive. In particular, our knowledge and understanding as to how the vertical transmission pathway of *Blattabacterium* is integrated into the postembryonic development of the host cockroaches are limited.

In this study, we conducted detailed histological and cytological observations of the localization of the bacteriocytes and the symbiotic bacteria during the postembryonic development of the German cockroach *Blattella germanica*. Based on these observations, we provide a hypothesis as to how the postembryonic developmental dynamics of the bacteriocytes are integrated into the vertical transmission and functioning of the endosymbiotic bacteria in cockroaches.

## Materials and methods

### Insect materials

A stock population of *B. germanica*, derived from around 100 individuals purchased from Sumika Technoservice Corporation, Takarazuka, Japan, was maintained in our laboratory at the National Institute of Advanced Industrial Science and Technology (AIST), Tsukuba, Japan. Developmental staging of nymphs was done as previously described [24, 72]. All insects were reared in plastic containers at 27°C under a 12 h light and 12 h dark regime in a climate chamber (MLR-352H-PJ, Panasonic) with insect feed (Insect Diet I, Oriental Yeast Co., Ltd.) and water.

### Fluorescence *in situ* hybridization (FISH)

The insects were dissected in a phosphate-buffered saline (PBS: 0.8% NaCl, 0.02% KCl, 0.115% Na_2_HPO_4_, 0.02% KH_2_PO_4_ [pH 7.4]). The isolated ovaries were fixed in Carnoy’s solution (ethanol: chloroform: acetic acid = 6: 3: 1) for about 45 min, and preserved in 70% ethanol in a refrigerator until use. The samples were incubated in PBT (PBS containing 0.1% Tween 20) for a few hours and then washed with 1 M Tris buffer (pH 9.0) several times until the color of the samples became translucent (approx. 3 h). This treatment was for the removal of accumulated uric acid crystals (primarily in the visceral fat bodies) as they disturb fluorescence imaging. FISH was conducted essentially as described previously [73] using a fluorochrome-labeled oligonucleotide probe Sul664R [5′-Alexa555-CCM CACATT CCA GYT ACT CC-3′] (100 pmol/ml) targeting 16S rRNA of *Blattabacterium* [74] with counterstaining by 4’,6-diamidino-2-phenylindole (DAPI) overnight. These samples were mounted in 90% glycerol and observed under a fluorescence stereomicroscope (M165FC, Leica, Germany) and confocal laser scanning microscope (AX R, Nikon, Japan).

### Cytological visualization of muscle fibers

The insects were dissected as described above and the isolated tissues were fixed in buffered PFA (PBS containing 4% paraformaldehyde) and thoroughly washed with PBT. After incubation in PFA, the tissue samples were stained with Alexa Fluor 488 phalloidin (Invitrogen, USA), washed thoroughly with PBT, and subjected to DNA staining with PBS supplemented with 1 μg/mL DAPI, and observed under a fluorescence stereomicroscope (M165FC, Leica, Germany) and confocal laser scanning microscope (AX R, Nikon, Japan).

### Electron microscopy

For backscattered electron scanning electron microscopy (BSE-SEM), ovaries were dissected from adult females in cold PBS, and individual ovarioles were separated using needles. The samples were fixed overnight at 4°C in the mixture of 4% paraformaldehyde (Cat. No. 162-16065, Wako Pure Chemical Industries) and 2% glutaraldehyde (Sigma-Aldrich, G5882) in PBS, incubated with 2% osmium tetroxide in distilled water at 4°C for 2 h, dehydrated in a graded ethanol series (50%, 60%, 70%, 80%, 90%, 95%, 99%, and 100%), treated with 100% propylene oxide followed by Epon 812, and embedded in Epon 812. The hardened Epon blocks were sectioned using an ultramicrotome (ARTOS 3D, Leica) equipped with a diamond knife (SYM jumbo, 45 degrees, SYNTEK) to obtain 240 nm serial sections. The serial sections were collected on cleaned silicon wafer strips held by a micromanipulator (MN-153, NARISHIGE). The sections were stained at room temperature with 2% (w/v) aqueous uranyl acetate for 20 min and Reynolds’ lead citrate for 3 min. Images were obtained by a scanning electron microscope (JSM-7900F, JEOL).

## Results

### Developmental stages of *B. germanica*

To investigate the post-embryonic developmental stages of *B. germanica*, first instar nymphs were sexed after hatching and maintained individually under a defined rearing condition (27 °C, 12 h light and 12 h dark, fed with insect feed) (Fig. 1a). These insects were checked everyday and the date of each molt was recorded. The insects took about 20 days to hatch, molted to second instar on day 5-6, third instar on around day 12, fourth instar on around day 18, fifth instar on day 28-30, sixth instar on day 35-40, and adults emerged on around day 40 (Fig. 1b). After eclosion, females took several days to reach sexual maturity [75], during which they were not receptive to copulation. We observed that, by day 10 post-eclosion, all females were receptive to copulation. Thus, we judged that the adult females became sexually mature by day 10 post-eclosion.

**Fig. 1.**
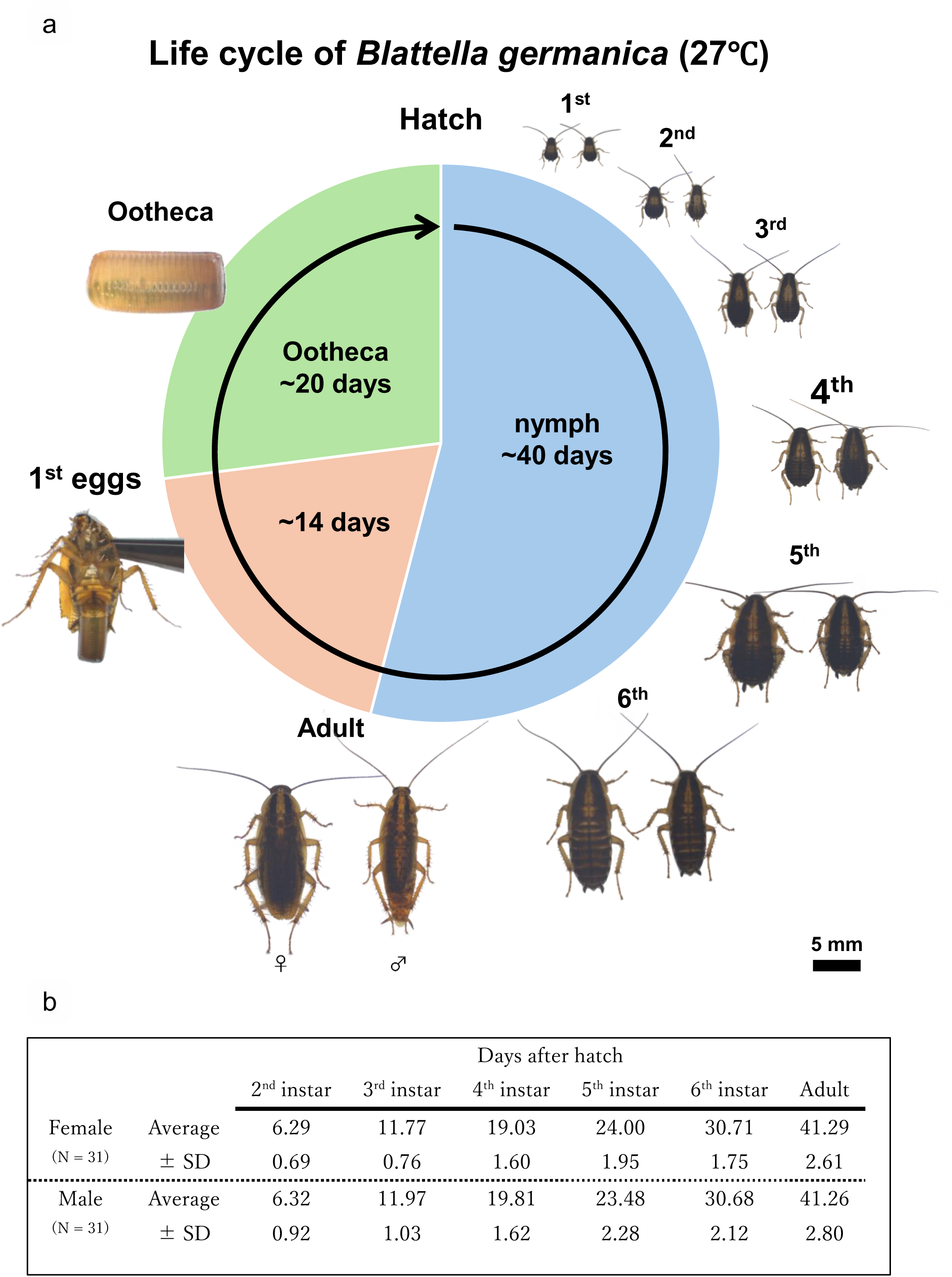
Life-cycle of the German cockroach *Blattella germanica* reared at 27℃. (**a**) Photos taken on the first day of each developmental stage (left: female, right: male). (**b**) Duration of the developmental stages of 62 insects (days, mean ± standard deviation).

### Post-embryonic ovary development of *B. germanica*

To investigate the post-embryonic developmental process of the ovaries, we dissected and observed the developing ovaries of the staged individuals as follows: early first instar nymphs (< 3 days post-hatch), late first instar nymphs (> 5 days post-hatch), second to sixth instar nymphs, sexually immature adults (< 3 days post eclosion), and sexually mature adults (> 10 days post eclosion). The ovaries of the insects in the early developmental stages were extremely small and underdeveloped, but their anatomical position was consistent, being located close to the third and fourth abdominal tergites. The ovaries were distinguishable due to their translucent appearance in contrast to the opaque white fat bodies surrounding them (Fig. 2a; Fig. 3a). These ovaries were subjected to morphological observations and FISH visualization of *Blattabacterium* and bacteriocytes.

**Fig. 2.**
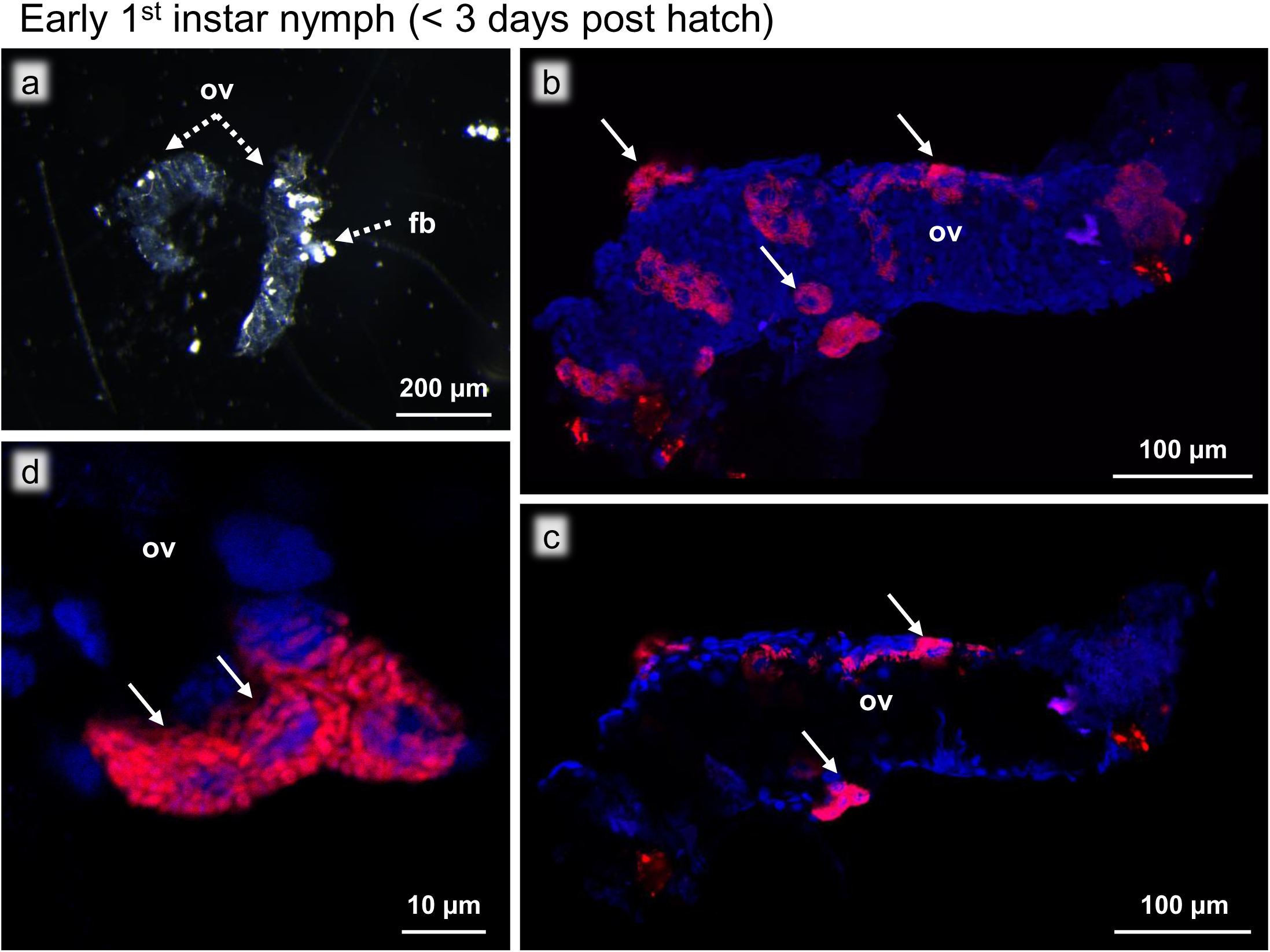
Ovaries of an early first instar nymph (< 3 days post hatch). (**a**) Bright field image of the ovaries. Translucent ovaries (ov) are associated opaque fat bodies (fb). (**b**) FISH image of the whole ovary. The bacteriocytes (arrows) full of *Blattabacterium* are visualized in red with DNA counterstaining in blue. (**c**) FISH optical section of the whole ovary. The bacteriocytes are not in the ovary but remaining on the surface of the ovary. (**d**) Magnified FISH image of the bacteriocytes on the ovarial surface. Red and blue indicate FISH signals of *Blattabacterium* 16S rRNA and DNA signals of DAPI staining, respectively.

**Fig. 3.**
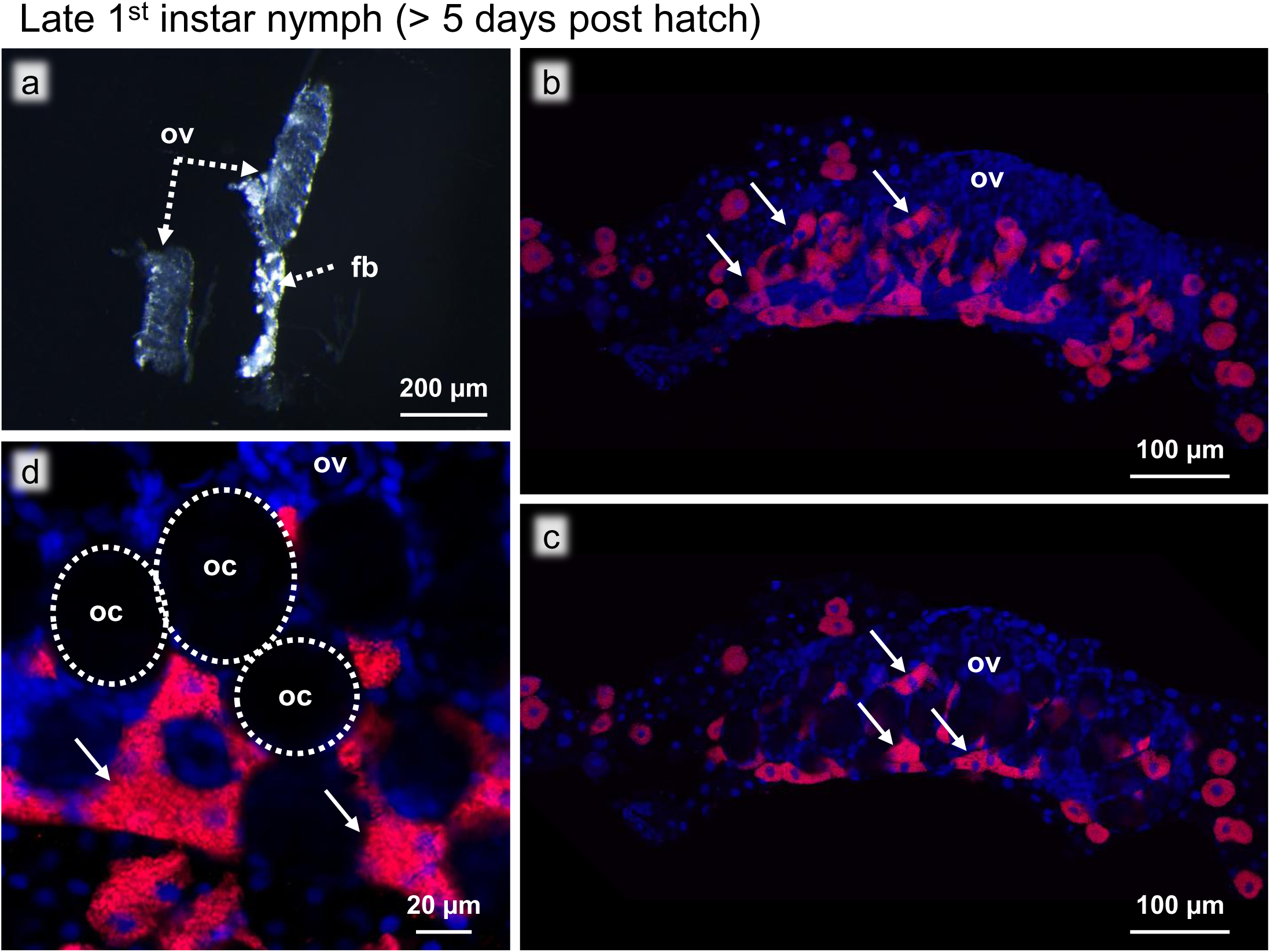
Ovaries of a late first instar nymph (> 5 days post hatch). (**a)** Bright field image of the ovaries. (**b**) FISH image of the whole ovary. (**c**) FISH optical section of the whole ovary. (**d**) Magnified FISH image of the oocytes and the intraovarial bacteriocytes. Dotted circles indicate the oocytes. Abbreviations: fb, fat body; oc, oocyte; ov, ovary. Arrows show bacteriocytes. Red and blue indicate FISH signals of *Blattabacterium* 16S rRNA and DNA signals of DAPI staining, respectively.

### Early first instar nymphs (< 3 days post-hatch)

The ovaries of early first instar nymphs were small (approx. 400 μm in length) and translucent, in which neither oocytes nor ovarioles were recognizable (Fig. 2a). FISH observations showed that the tiny ovaries were still uninfected with *Blattabacterium*, while multiple bacteriocytes were found on the surface of the ovaries (Fig. 2b). The FISH optical sections confirmed that the bacteriocytes were not inside but on the surface of the ovaries (Fig. 2c, d).

### Late first instar nymphs (> 5 days post-hatch)

The morphological characteristics of the ovaries of late first instar nymphs remained largely the same, but the developing oocytes became recognizable (Fig. 3a-c). The FISH images revealed a drastic increase in the number of bacteriocytes around and on the surface of the ovaries compared to the early first instar nymphs (Fig. 3b). In addition, the FISH optical sections showed that many bacteriocytes were located within the ovaries in the spaces between the oocytes (Fig. 3c). At this stage, *Blattabacterium* signals were restricted to the intraovarial bacteriocytes and the oocytes were still uninfected (Fig. 3d).

### Second instar nymphs

The morphological characteristics of the ovaries of second instar nymphs exhibited more adult-like traits compared to those of the prior stages, with distinctly recognizable oocytes and ovarioles (Fig. 4a), which were also clearly visible in the FISH images (Fig. 4b-c). The FISH images revealed that the bacteriocytes were no longer on the surface of the ovaries but in the intraovarial spaces (Fig. 4b, c). At this stage, notably, we observed that the surface of the oocytes was infected with *Blattabacterium*, where some images seemed to represent the process of bacteriocyte-to-oocyte symbiont transmission (Fig. 4d).

**Fig. 4.**
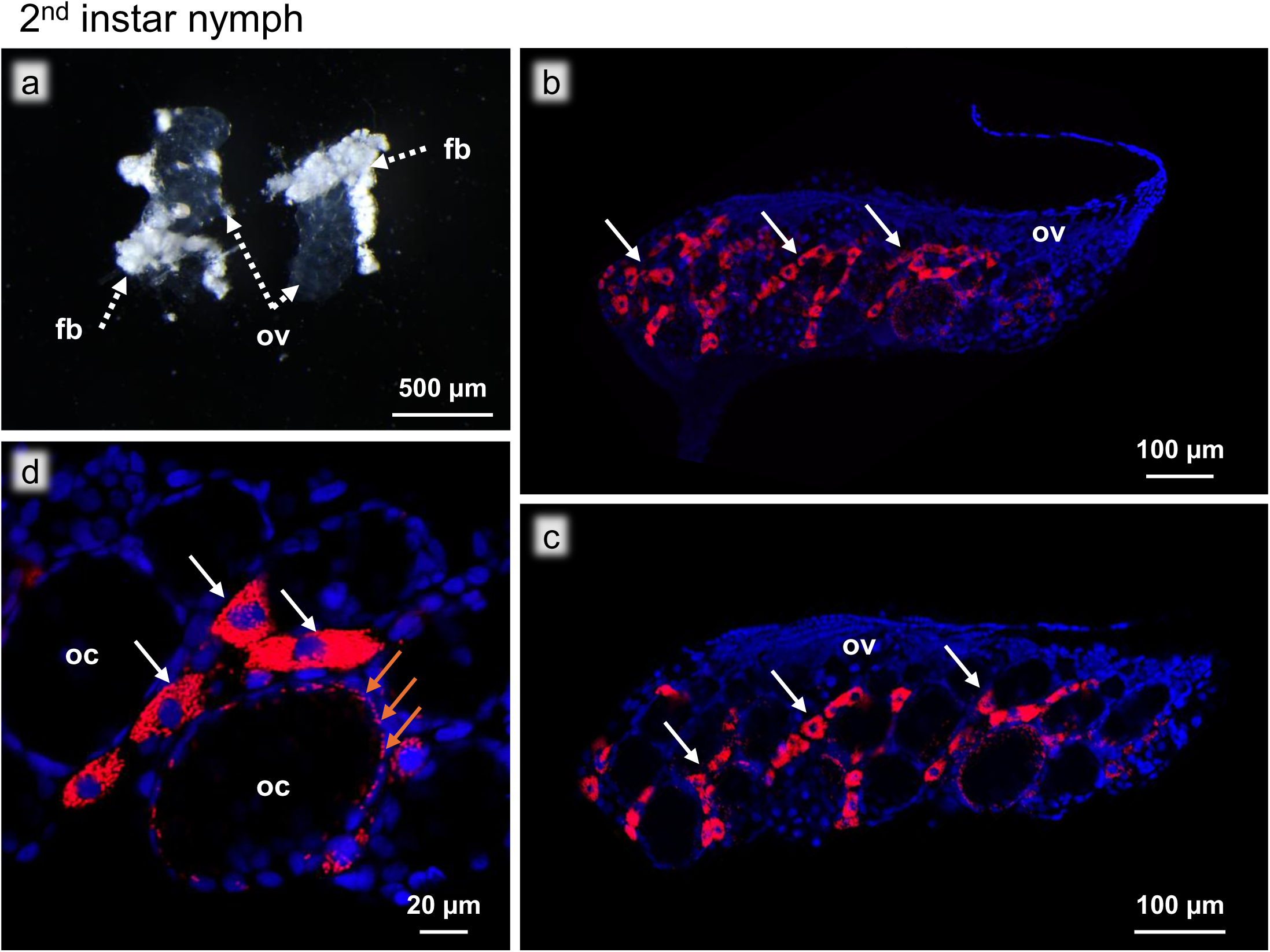
Ovaries of a second instar nymph. (**a**) Bright field image of the ovaries. (**b**) FISH image of the whole ovary. (**c**) FISH optical section of the whole ovary. (**d**) Magnified FISH image of the oocytes and the intraovarial bacteriocytes. Abbreviations: fb, fat body; oc, oocyte; ov, ovary. White arrows show bacteriocytes. Brown arrows show *Blattabacterium* on the oocyte surface. Red and blue indicate FISH signals of *Blattabacterium* 16S rRNA and DNA signals of DAPI staining, respectively.

### Third to sixth instar nymphs

The development of the ovaries from third to sixth instar nymphs was characterized by enlargement of the oocytes, elongation of the ovarioles, thickening of the oviducts, and overall increase in the ovarial size (Fig. 5a-d). FISH images revealed cytological characteristics similar to those of the second instar nymphs: the bacteriocytes are not on the ovary surface but in the intraovarial spaces, and the oocytes were infected with *Blattabacterium* (Fig. 5e-h). The sectioned FISH images showed that the bacteriocyte-to-oocyte transmission of *Blattabacterium* occurred throughout the nymphal stages (Fig. 5i-l). Notably, as the ovarial development proceeded, the intraovarial bacteriocytes, which were evenly distributed throughout the ovaries in the second instar nymphs, became concentrated towards the anterior end of the ovaries and the terminal filaments.

**Fig. 5.**
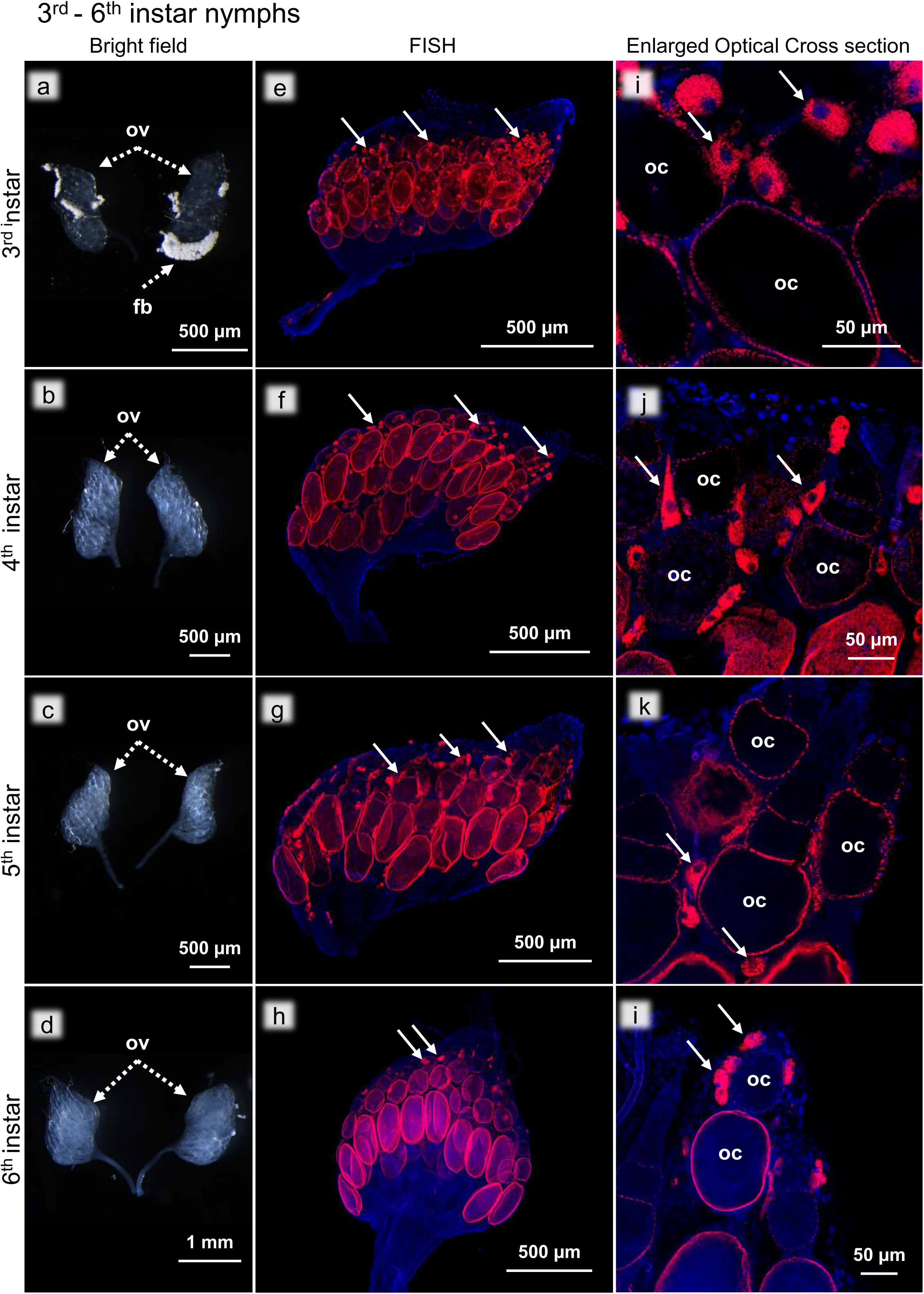
Ovaries of third to sixth instar nymphs. (**a-d**) Bright field images of the ovaries. (**e-h**) FISH images of the whole ovaries. (**i-l**) Magnified FISH images of the oocytes and the intraovarial bacteriocytes. (**a, e, i**) Third instar. (**b, d, j**) Fourth instar. (**c, f, k**) Fifth instar. (**d, h, l**) Sixth instar. Abbreviations: fb, fat body; oc, oocyte; ov, ovary. Arrows indicate bacteriocytes. Red and blue indicate FISH signals of *Blattabacterium* 16S rRNA and DNA signals of DAPI staining, respectively.

### Sexually immature adults (< 3 days post eclosion)

The ovaries of sexually immature adults were, except for less developed oocytes, morphologically similar to the ovaries of sexually mature adults (Fig. 6a; Fig. 7a). FISH observations showed that the oocytes were still infected with *Blattabacterium* (Fig. 6b). In the adult ovaries, it was evident that the ovarioles were bundled together by an outer membrane, so-called the peritoneal sheath, consisting of a layer of cells visible with fluorescent staining (Fig. 6b), which was not apparent in the nymphal ovaries. Fluorescent visualization of actin filaments using phalloidin revealed this membrane to be a net of actin filaments that surrounded the ovarioles (Fig. S1), which was also present around the ovaries of the sexually mature adults (Fig. S2), but not recognizable during the nymphal stages (Fig. S3). FISH images also showed that, at this stage, there were no intraovarial bacteriocytes but only bacteriocytes on the surface of the ovaries (Fig. 6c). Magnified FISH images revealed that these bacteriocytes were no longer in direct contact with the oocytes, where no bacteriocyte-to-oocyte transmission of *Blattabacterium* was seen (Fig. 6d). In addition, we were unable to identify any other bacteriocytes closely associated with the ovaries, nor any *Blattabacterium* being transmitted through another pathway (e.g. through the oviduct).

**Fig. 6.**
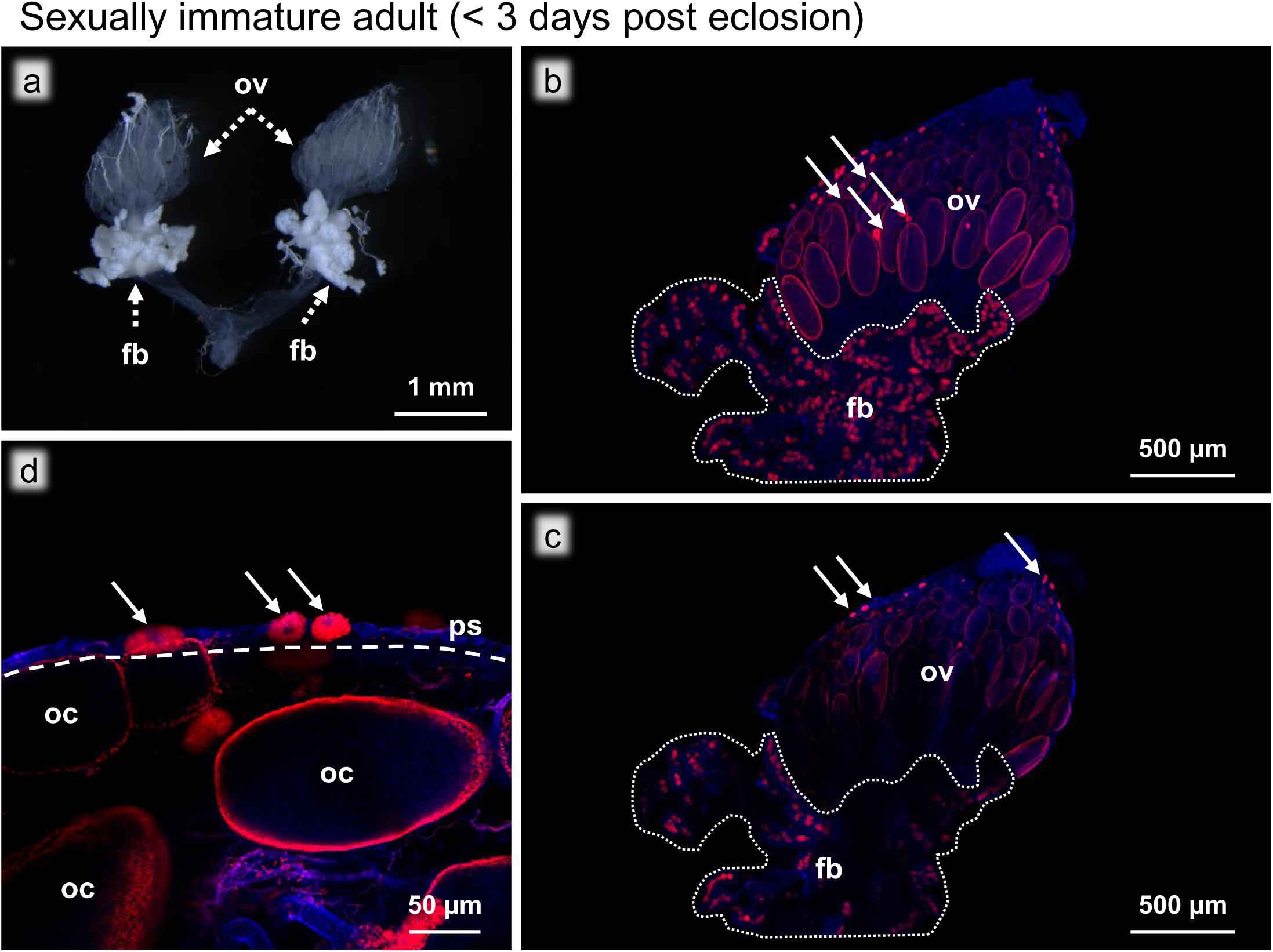
Ovaries of a sexually immature adult (<3 days post eclosion). (**a**) Bright field image of the ovaries. (**b**) FISH image of the whole ovary. (**c**) FISH optical section of the whole ovary. (**d**) Magnified FISH image of the oocytes and the bacteriocytes on the ovarial surface. The dashed line indicates the outer membrane (peritoneal sheath). The dotted area indicates the fat bodies. Abbreviations: fb, fat body; oc, oocyte; ov, ovary; ps, peritoneal sheath. Arrows show bacteriocytes. Red and blue indicate FISH signals of *Blattabacterium* 16S rRNA and DNA signals of DAPI staining, respectively.

**Fig. 7.**
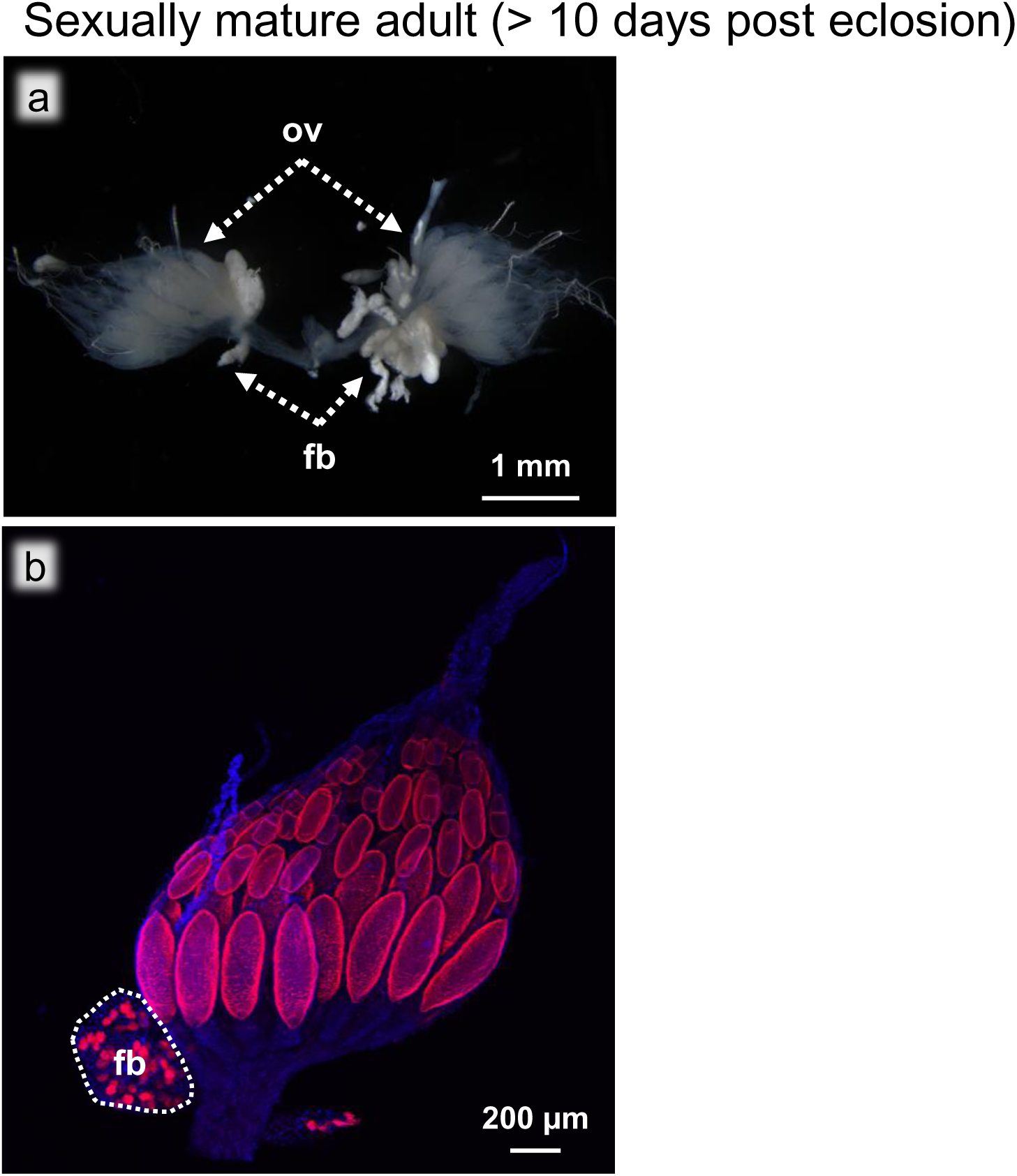
Ovaries of a sexually mature adult (>7 days post eclosion). (**a**) Bright field image of the ovaries. (**b**) FISH image of the whole ovary. Abbreviations: fb, fat body; ov, ovary. The dotted area indicates the fat bodies. Red and blue indicate FISH signals of *Blattabacterium* 16S rRNA and DNA signals of DAPI staining, respectively.

### Sexually mature adults (> 10 days post eclosion)

The most posterior oocytes of sexually mature adults were significantly larger and more developed, and were no longer translucent due to the accumulation of yolk granules (Fig. 7a). FISH revealed that the oocytes were still infected with *Blattabacterium*, but neither intraovarial bacteriocytes nor bacteriocytes on the ovarial surface were not found (Fig. 7b). As observed in the sexually immature adult insects, we were unable to observe any other transmission routes for the symbiotic bacteria, suggesting that no *Blattabacterium* cells are transmitted to the oocytes at this stage.

### FISH and ultrastructure of sexually mature ovarioles

Our observations suggested that, in *B. germanica*, the bacteriocyte-to-oocyte transmission of *Blattabacterium* only occurs from the second to sixth instar nymphal stages. However, considering that adult females of *B. germanica* can produce up to nine oothecae, each containing about 40 eggs, throughout their lifetime of 100-250 days [74], a question arises: How do the newly-formed oocytes for egg/ootheca production get infected with *Blattabacterium*? FISH observations of the ovarioles dissected from sexually mature adults revealed that, while the surface of the larger oocytes was all infected with *Blattabacterium* (Fig. 8a), the anterior part of the ovarioles was devoid of *Blattabacterium* infection (Fig. 8b). In order to gain insight into the route of *Blattabacterium* infection to oocytes, we conducted ultrastructural observations of the ovarioles dissected from sexually mature adults using electron microscopy. Hereafter, we adopted the names of the ovariole regions Zone I to Zone V (Fig. 8; Fig. 9a) after a previous detailed study on oogenesis of the American cockroach *Periplaneta americana* [76].

**Fig. 8.**
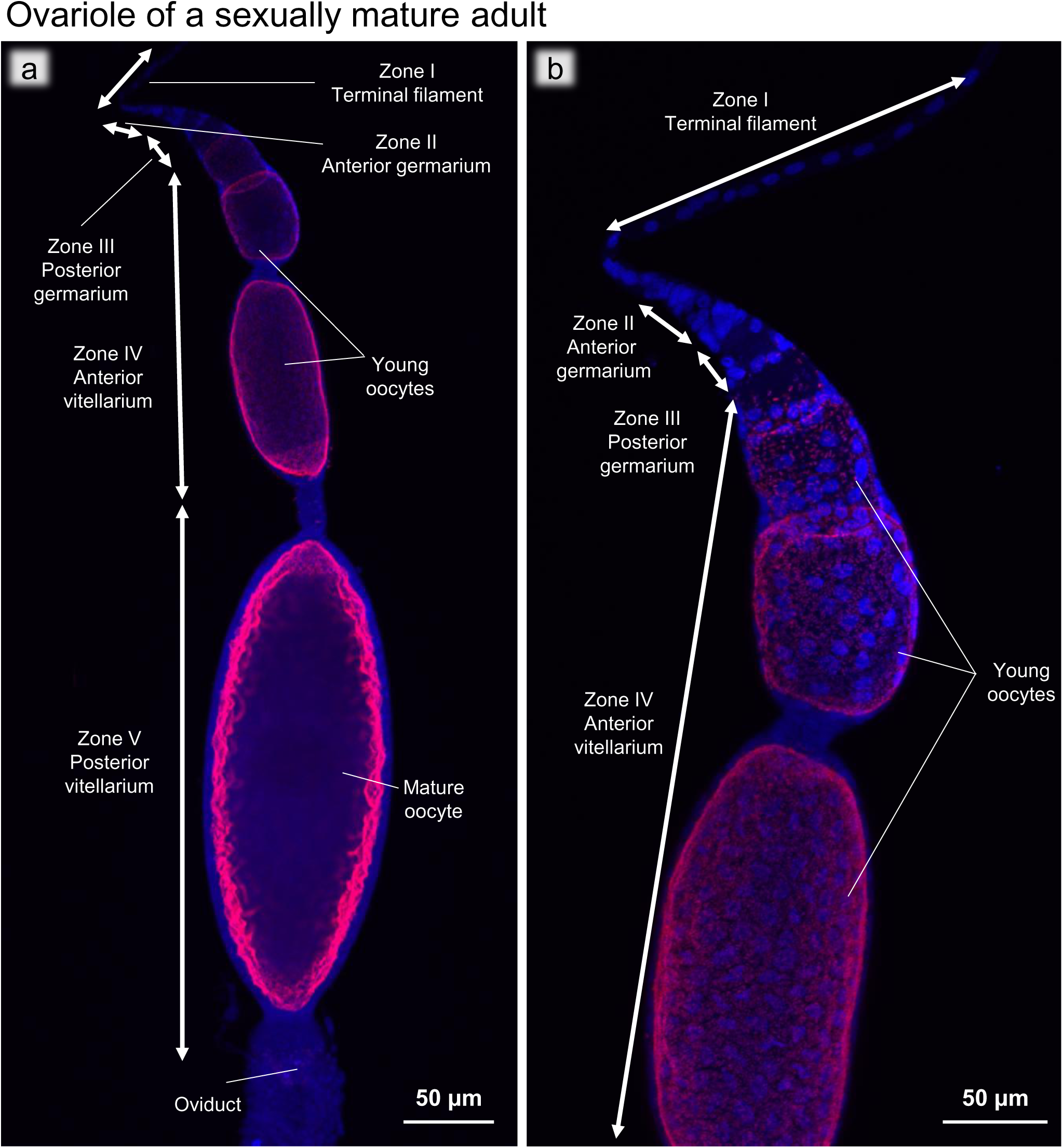
FISH images of an ovariole of a sexually mature adult. (**a**) FISH optical section of the whole ovariole. (**b**) Magnified FISH image of the anterior end of the ovariole. Red and blue indicate FISH signals of *Blattabacterium* 16S rRNA and DNA signals of DAPI staining, respectively.

**Fig. 9.**
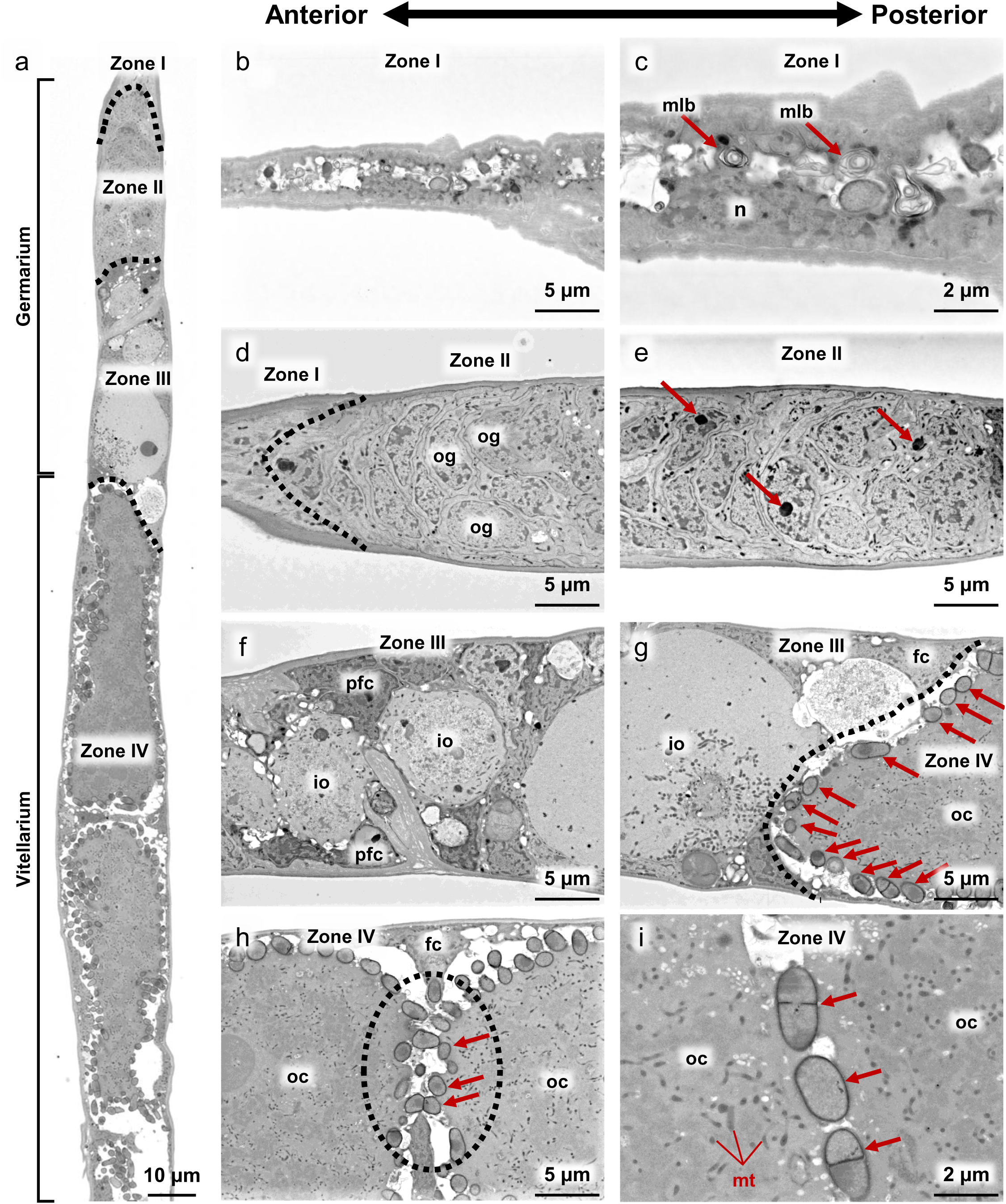
Electron microscopic images of the ovariole of a sexually mature adult. (**a**) BSE-SEM longitudinal section image of the tip region of the ovariole, which depict the zones I, II, III and IV defined by a previous study [76]. (**b, c**) Zone I, the terminal filament. Arrows indicate multilamellar bodies. (**d**) Zone I-Zone II, the terminal filament-germarium boundary indicated by a dotted line. (**e**) Zone II, the mid region of the germarium. Arrows indicate electron dense bodies in differentiating oogonia. (**f**) Zone III, the anterior region of the germarium, in which immature primary oocytes develop and increase in size. (**g**) Zone III-Zone IV, the germarium-vitellarium boundary indicated by a dotted line. In the anterior part of the vitellarium, young oocytes are arranged linearly, accumulating yolk and mitochondria in the ooplasm, and surrounded by follicle cells. On the surface of the young oocytes, many *Blattabacterium* cells (arrows) are seen. (**h**) Zone IV, the anterior region of the vitellarium, where a dotted circle highlights *Blattabacterium* cells located between and contacting two young oocytes. (**i**) Zone IV, the magnified image of the *Blattabacterium* cells bridging two young oocytes. Note that some *Blattabacterium* cells are in division with septum [91]. Numerous mitochondria are seen in the ooplasm. Abbreviations: fc, follicle cell; io, immature oocyte; mlb, multilamellar body; n, nucleus; io, immature oocyte; oc, oocyte; og, oogonium; pfc, prefollicular cells; yo, young oocyte.

#### Zone I

The terminal filament, a stack of flattened somatic cells, acts as a physical anchor that attaches to the dorsal cuticle, keeps the ovaries suspended, and has no cells associated with oogenesis [77]. The terminal filament cells were characterized by degenerative cytology with multilamellar bodies within the cytoplasm (Fig. 9b, c).

#### Zone II

In the germarium region (Fig. 9a, d, e), the germline stem cells differentiate into oogonia and develop into primary oocytes [78]. In the anterior germarium region, undifferentiated cells with a large nucleus were densely seen, which probably represented proliferating germline stem cells and oogonia (Fig. 9d, e). In the posterior part of Zone II, some cells contained an electron dense body in the nucleus, which may represent primary oocytes (Fig. 9e).

#### Zone III

In the posterior germarium region, immature oocytes and prefollicular cells differentiated (Fig. 9f) and follicle cells became distinguishable. The immature oocytes were arranged non-linearly and drastically increased in size toward the posterior part, developing a large nucleus and beginning to accumulate mitochondria in the cytoplasm (Fig. 9f, g). *Blattabacterium* cells were observed at the posterior end of Zone III, being in contact with Zone IV (Fig. 9g), which was concordant with the FISH visualization patterns of *Blattabacterium* in the ovarioles (Fig. 8).

#### Zone IV

In the anterior vitellarium region, young oocytes were linearly arranged, increasing in size, surrounded by a follicle cell layer, and accumulating mitochondria in the ooplasm (Fig. 9g, h). In Zone IV, numerous *Blattabacterium* cells were found in the space between the young oocyte and the follicle cell layer, indicating the major site of *Blattabacterium* localization in the ovarioles (Fig. 9g, h) and confirming the FISH visualization patterns of *Blattabacterium* in the ovarioles (Fig. 8). In Zone IV, notably, when we observed neighboring young oocytes, the follicle cell layers surrounding the oocytes occasionally exhibited breaks, where *Blattabacterium* cells were located between and contacting with neighboring young oocytes (Fig. 9h, i), suggesting that the symbiotic bacteria can move across and infect to neighboring young oocytes through this passage within each ovariole. These passages were only observed in the center of the ovarioles, and the oocytes were completely separated from each other by follicle cells in the periphery of the ovariole (Fig. S4).

#### Zone V

In the posterior vitellarium region, the oocytes accumulated yolk granules, drastically increased in size, and formed chorion on the surface, thereby turning into mature oocytes ready for oviposition (Fig. 8a; BSE-SEM image not shown).

## Discussion

In this study, we present a detailed and comprehensive description of the vertical transmission pathway of *Blattabacterium* in *B. germanica* during the post-embryonic development, as schematically summarized in Fig 10. Our study identified three distinct stages of vertical symbiont transmission with dynamic bacteriocyte migration: (1) the pre-infection phase, where surrounding bacteriocytes move into the developing ovaries that are initially devoid of bacteriocytes (Fig 10a,b), (2) the infection phase, where *Blattabacterium* cells from the intraovarial bacteriocytes are transmitted to the developing oocytes (Fig 10c,d), and (3) the post-infection phase, where the intraovarial bacteriocytes are lost and bacteriocyte-to-oocyte infection no longer occurs (Fig 10e, f). In addition, we observed that, in the ovarioles of sexually mature adult insects, the germline cells in the germarium are uninfected and *Blattabacterium* seems to be transmitted from mature oocyte to young oocyte within each of the ovarioles.

**Fig. 10.**
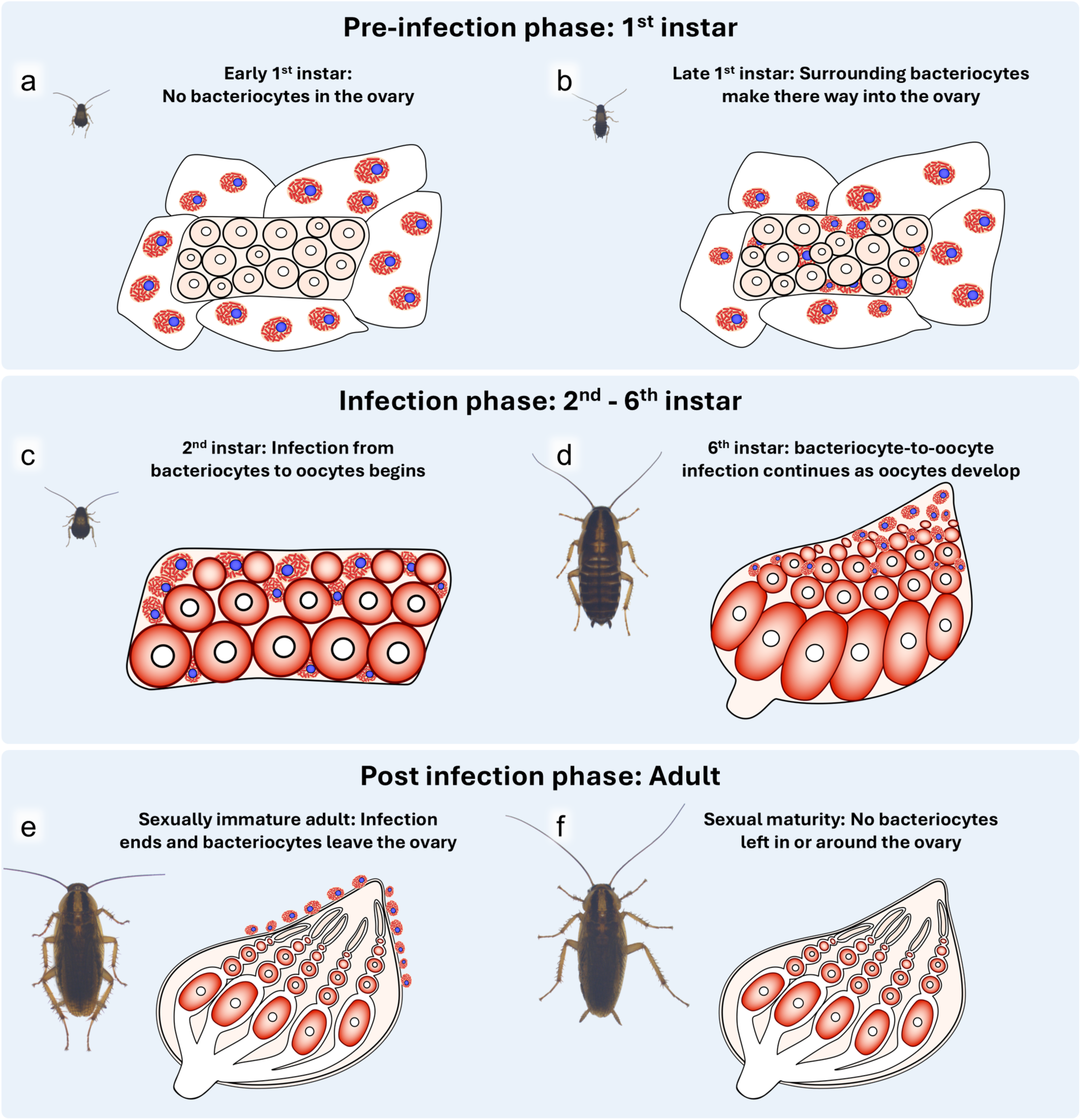
Schematic diagram of the ovarial morphogenesis, bacteriocyte migration, and symbiont transmission during the post-embryonic development of *B. germanica*. (**a**) Early first instar. (**b**) Late first instar. (**c**) Second instar (**d**) Sixth instar. (**e**) Sexually immature adult. (**f**) Sexually mature adult. See text for detail.

The pre-infection phase is characterized by the surrounding bacteriocytes migrating into the ovaries during the first instar stage. Just after hatching, the developing ovaries are uninfected with *Blattabacterium* and only a few bacteriocytes adhere on the ovarial surface. This configuration changes soon, as observed in the late first instar, wherein the number of bacteriocytes around the developing ovaries rapidly increases. In addition, notably, many of these bacteriocytes get entry into the ovaries and occupy the spaces between the developing oocytes. Despite the intraovarial localization of the bacteriocytes harboring *Blattabacterium*, the oocytes are still uninfected at this stage and *Blattabacterium* cells are still confined within the intraovarial bacteriocytes. Although speculative, we believe that these bacteriocytes migrated from surrounding fat bodies, as we have never observed any bacteriocytes on the surface of the ovaries undergoing cell division.

The infection phase is characterized by the bacteriocyte-to-oocyte transmission of *Blattabacterium* cells. In the ovaries of second instar nymphs, no bacteriocytes are found on the surface of the ovaries, and all visible bacteriocytes are located in intraovarial spaces. At this stage, *Blattabacterium* cells are transmitted from the intraovarial bacteriocytes to their neighboring oocytes. It should be noted that the image shown (Fig. 4d) was taken from an individual only a day after the first molt, and yet the symbiont transmission had already started, suggesting that the infection had likely started immediately after molting. Bacteriocyte-to-oocyte symbiont transmission continues throughout the rest of the nymphal stages. As the ovaries develop through the third to sixth instar stages, the intraovarial bacteriocytes, which are evenly distributed throughout the ovary during the second and third instar stages, become more concentrated towards the anterior region of the ovaries. Whether these developmental patterns are manifested by movement of the bacteriocytes within the ovaries or caused by the growth of the surrounding ovarial cells is unclear.

The post-infection phase is characterized by the intraovarial bacteriocytes exiting the ovaries and the termination of the bacteriocyte-to-oocyte symbiont transmission. Even after adult molting, some bacteriocytes are associated with the ovaries. However, close inspection revealed that these bacteriocytes are all on the outer surface of the ovaries and intraovarial bacteriocytes are no longer found, in which no bacteriocyte-to-oocyte symbiont transmission is observed. These bacteriocytes associated with the ovarial surface are completely absent in the ovaries of sexually mature adults. In addition, we were unable to observe any other migration routes of *Blattabacterium* (ex. through the oviduct, follicle cells, etc.), suggesting that bacteriocyte-to-oocyte transmission of *Blattabacterium* is terminated upon adult emergence of *B. germanica*.

On the adult ovaries, we found a membranous structure that encases the whole ovarioles each side together. Phalloidin staining revealed that this structure is at least partially composed of actin filaments that cover the ovarioles like a net, which is likely to represent a structure called the peritoneal sheath described from the ovaries of diverse insects [79–82]. While biological functions of the peritoneal sheath have been suggested as keeping the integrity of the ovaries and helping oviposition by muscular contraction [80], the potential involvement of the structure in the process of the bacteriocyte migration/exclusion in cockroaches is of interest.

The close association of the cockroach bacteriocytes with nymphal ovaries has been reported in previous studies [42, 43, 48, 53, 55]. However, most of these studies did not focus on the infection dynamics during postembryonic development including adult stages. Consequently, these previous studies failed to recognize the fact that bacteriocyte-to-ovary symbiont transmission no longer proceeds at the adult stages even though there are many bacteriocytes in the surrounding fat bodies. Our study provides the first detailed and comprehensive morphological and histological observations on the infection dynamics and transmission pathway of the *Blattabacterium* endosymbiont during the postembryonic development of the cockroach host.

Generally, bacteriocytes and bacteriomes exhibit strict localization [24–26] and postembryonic movement of bacteriocytes is uncommon. On the other hand, bacteriocytes/bacteriomes are often closely associated with the ovaries [16], where specialized tissues/organs for facilitating vertical symbiont transmission are often found and symbiont migration occurs there [83–87]. Notably, another case of postembryonic bacteriocyte migration has been reported in whiteflies, in which the maternal bacteriocytes migrate directly into each egg and vertical symbiont transmission establishes thereby [88, 89]. The molecular and cellular mechanisms underlying the bacteriocyte migration during the host insect development are totally unknown and deserve future studies.

## Conclusion

Here, we described the vertical transmission processes of the *Blattacterium* endosymbiont through ovarial infection during the post-embryonic development of the ovaries and the ultrastructure of the ovarioles in sexually mature females of *B. germanica*. We uncovered that there are three distinct phases of vertical symbiont transmission and provided the evidence of a previously unrecognized pathway of vertical symbiont transmission. This study represents the most up-to-date and comprehensive description of the vertical transmission pathway of *Blattabacterium*. This information is not only beneficial in our study of the cockroach-*Blattabacterium* symbiotic system but also aids in our understanding of the evolutionary adaptations required to realize intracellular microbial symbiosis.

## Acknowledgments

We thank Haruyasu Kohda and Keiko Okamoto-Furuta for their support in electron microscopic analysis, Cassandra Lord for schematic illustrations, and Kohei Oguchi for helpful comments on the ovarial development of blattodean insects.

## Contributions

T.N. conceived the research, reared all insects, processed the samples for microscopic observations, and conducted fluorescent microscopy. M.M. assisted with fluorescence microscopy and preparation of BSE-SEM samples, and supervised experiments. T.H. and T.K. performed electron microscopy. T.N. and T.F. wrote the manuscript. All authors contributed to and approved the final version of the manuscript.

## Ethics declarations

### Ethics approval and consent to participate

Not applicable.

## Consent for publication

Not applicable.

## Supplementary materials

**Fig. S1.**
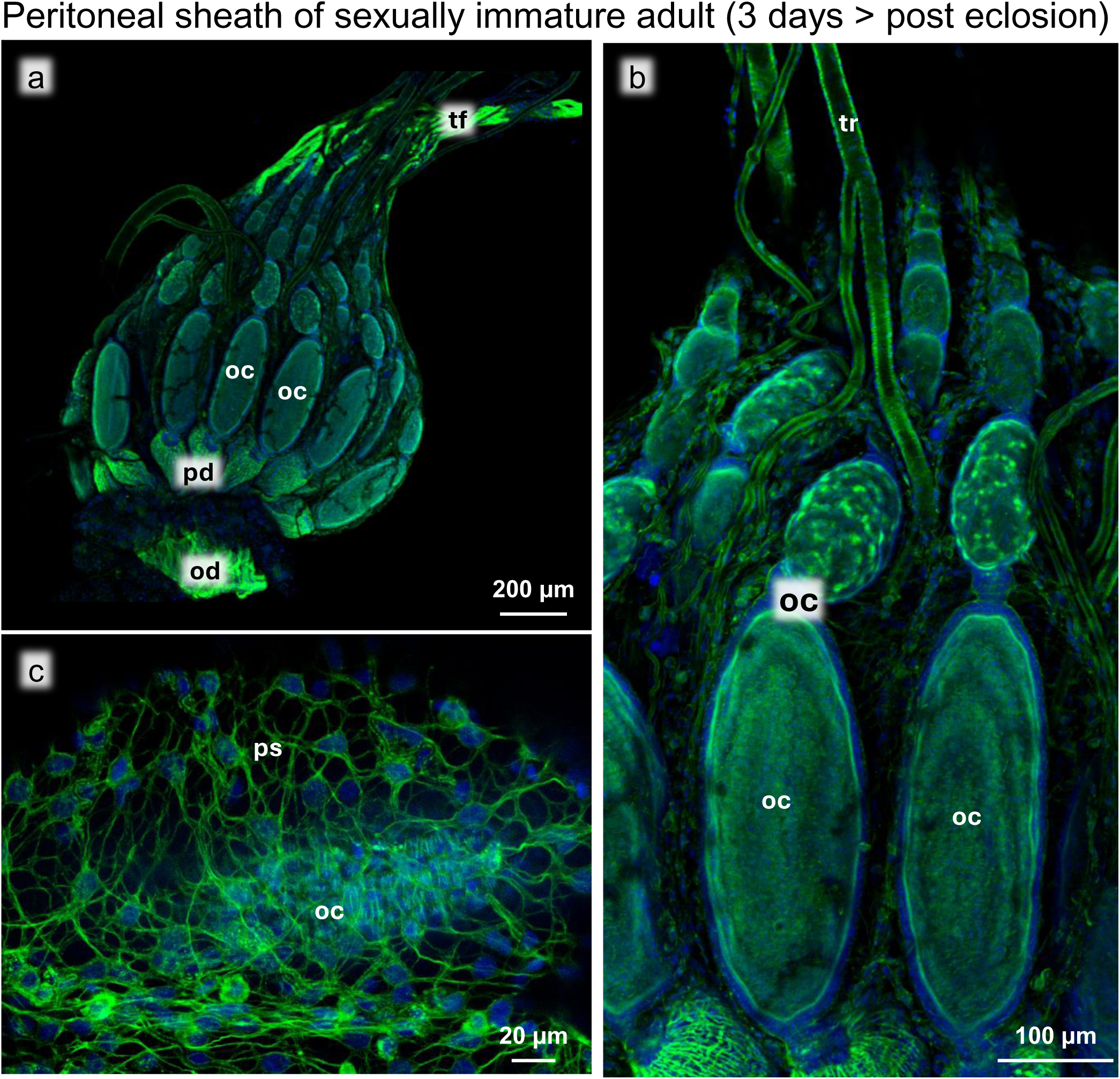
Actin-stained ovary of a sexually immature adult. (**a**) Image of the whole ovary. (**b**) Magnified image of the ovarial surface. (**c)** Magnified image of the outer membrane (peritoneal sheath). Abbreviations: oc, oocyte; od, oviduct; pd, ovarial pedicle; ps, peritoneal sheath; tf, terminal filament; tr, trachea.

**Fig. S2.**
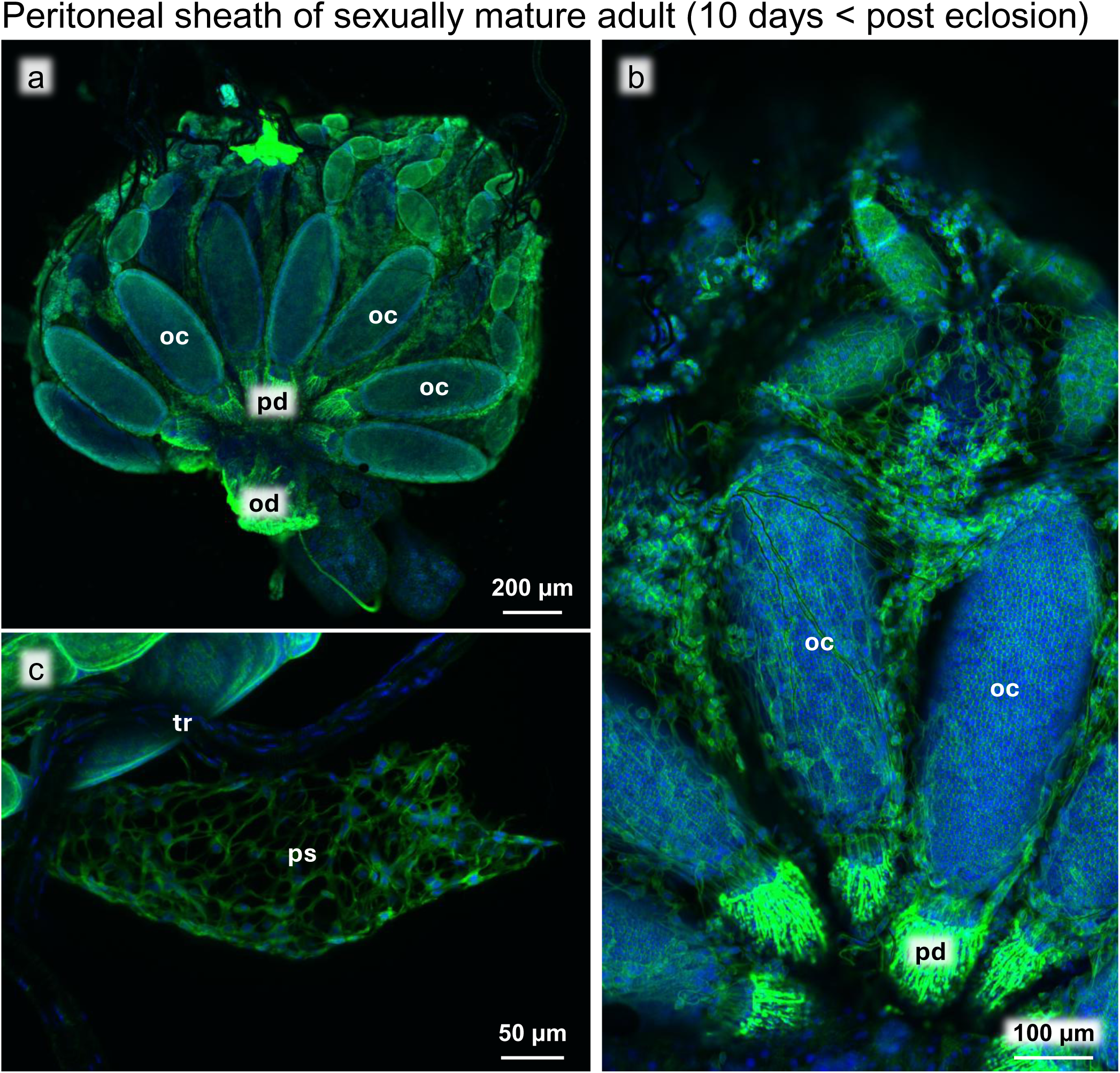
Actin-stained ovary of a sexually mature adult. (**a)** Image of the whole ovary. (**b)** Magnified image of the ovarial surface. (**c)** Image of the partially removed outer membrane (peritoneal sheath). Abbreviations: oc, oocyte; od, oviduct; pd, ovarial pedicle; ps, peritoneal sheath; tr, trachea.

**Fig. S3.**
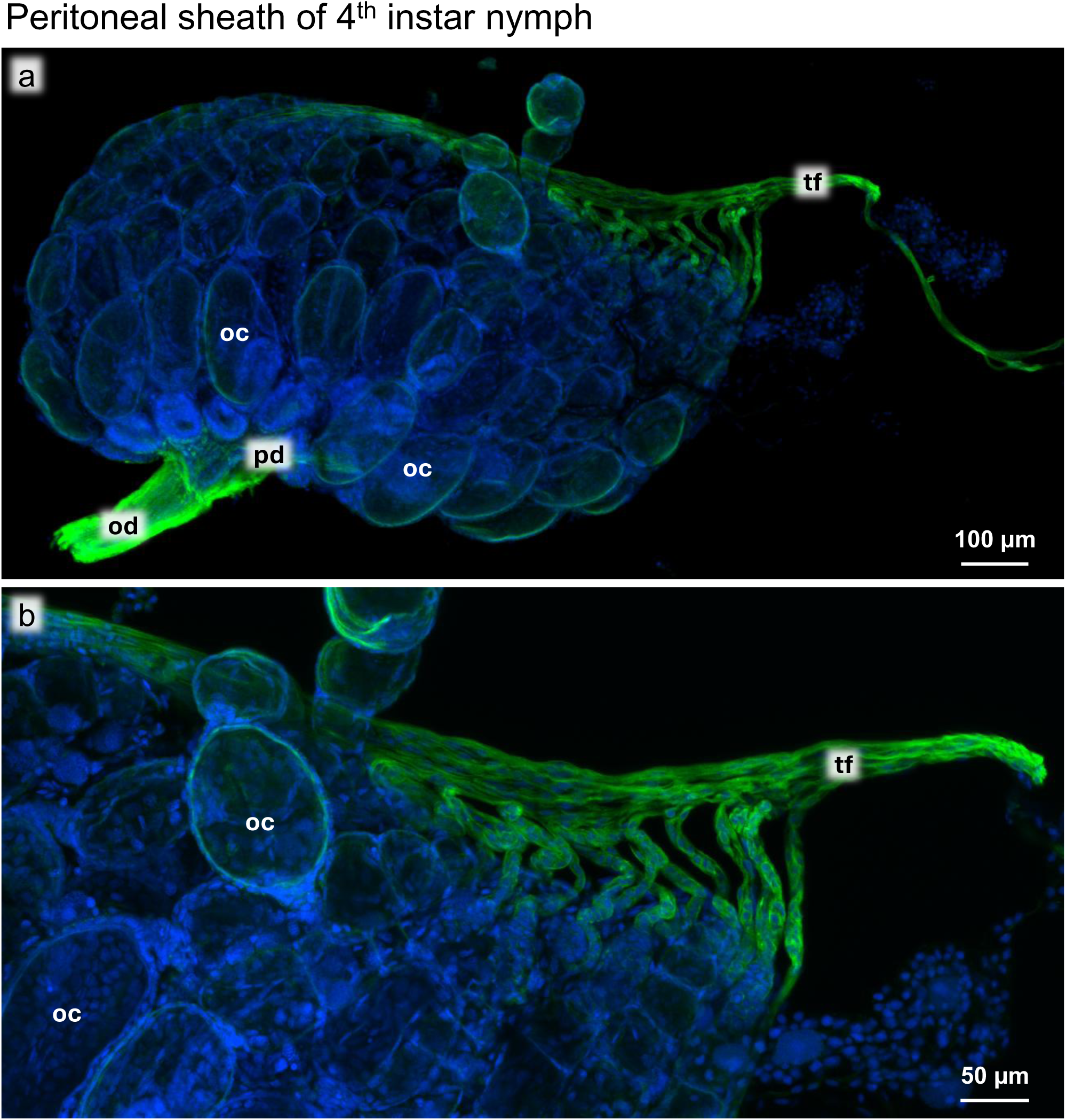
Actin-stained ovary of a fourth instar nymph. (**a**) Image of the whole ovary. (**b**) Magnified image of the ovarial surface. Abbreviations: oc, oocyte; od, oviduct; pd, ovarial pedicle; tf, terminal filament.

**Fig. S4.**
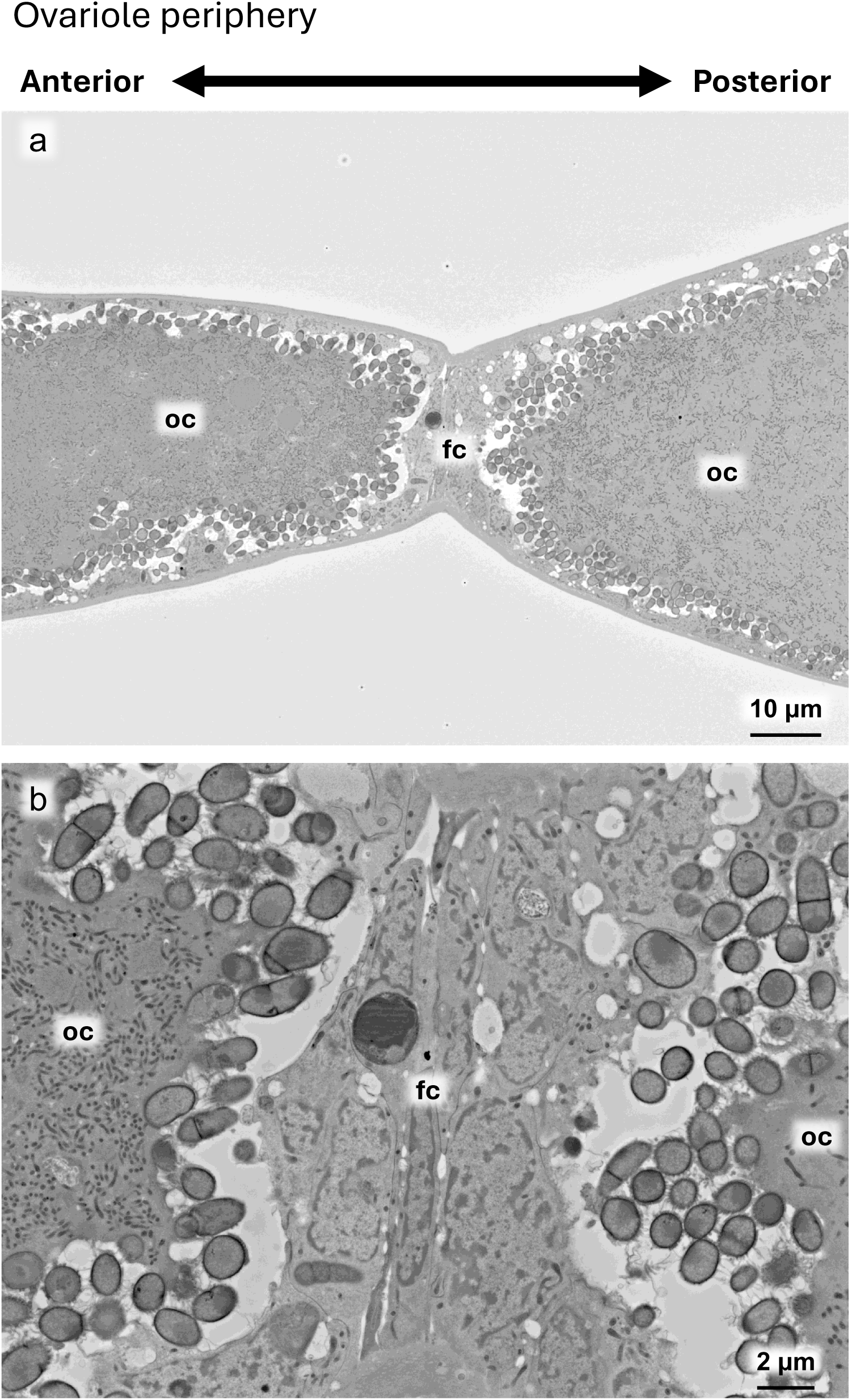
Electron microscopic images of Zone IV of the ovariole periphery. (**a**) Wider view of the follicle cell layer separating the two oocytes. (**b**) Magnified image of the follicle cell layer. Abbreviations: oc, oocyte; od, oviduct; pd, ovarial pedicle; tf, terminal filament.

